# Transcriptome-wide analysis suggests piRNAs preferentially recognize the coding region of mRNAs in *C. elegans*

**DOI:** 10.1101/2022.06.08.495319

**Authors:** Wei-Sheng Wu, Jordan S. Brown, Sheng-Cian Shiue, Dong-En Lee, Donglei Zhang, Heng-Chi Lee

## Abstract

**Background:** PIWI-interacting RNAs (piRNAs) protect genome integrity by silencing transposon mRNAs and some endogenous mRNAs in various animals. However, *C. elegans* piRNAs fail to trigger gene silencing at many sequence-based predicted targeting sites.

**Results:** To gain insights into the mechanisms that control piRNA silencing capability, we compared the transcriptome-wide predicted piRNA targeting sites to the *in vivo* piRNA binding sites. Surprisingly, while predicted piRNA targeting sites are enriched in 3’ UTRs, we found that *C. elegans* piRNAs preferentially bind to coding regions (CDS) of target mRNAs, leading to preferential production of secondary silencing small RNAs in the CDS. Furthermore, our analyses suggest that Argonaute protein CSR-1 protects mRNAs from piRNA silencing through two distinct mechanisms – by inhibiting piRNA binding across the entire CSR-1 targeted transcript, and by inhibiting secondary silencing small RNA production locally at CSR-1 bound sites. However, CSR-1 is not responsible for the piRNA binding preference for the CDS.

**Conclusions:** Our work identifies the CDS as the critical region that is uniquely competent for piRNA silencing in *C. elegans*. We speculate that the preference for CDS recognition by piRNAs may represent a mechanism to counteract the evolution of foreign protein-coding RNAs that evade piRNA surveillance.

## Background

Small non-coding RNAs, such as miRNAs and piRNAs, play important roles in gene regulation by guiding Argonaute proteins to target RNAs with sequence complementarity (1). miRNAs play critical roles in various developmental processes by regulating the expression of endogenous genes (2). PIWI-interacting RNAs (piRNAs) are germline-enriched small RNAs found in diverse animals that silence non-self nucleic acids, such as transposons and viruses (3–7). One unique characteristic of piRNAs is their sequence diversity - tens of thousands of sequence-distinct piRNAs are produced. To identify mRNA targets regulated by these diverse piRNAs, both sequence-based approaches as well as experimental approaches have been reported; using bioinformatic analyses and *in vivo* reporter assays, previous studies have identified the piRNA targeting rules in *C. elegans* (8,9). In *C. elegans*, piRNAs identify their targets through base pairing, requiring nearly perfect complementarity at seed regions and tolerating several mismatches in the remaining sequence. Several predicted piRNA targeting sites have been confirmed to play a regulatory role *in vivo*, and removal of predicted piRNA targeting sites allows for stable expression of several silencing-prone transgenes (8,9). piRNAs are predicted to bind various germline-silenced mRNAs, and surprisingly, many germline-expressed mRNAs as well (8,10). CLASH (crosslinking, ligation and sequencing of hybrids) is a powerful biochemical approach to identify *in vivo* small RNA binding sites (11); protein-bound small RNAs are ligated to their target mRNAs and these hybrid molecules, comprised of small RNAs and mRNA fragments, are sequenced to identify putative direct interactions. A recent study has applied CLASH analyses of *C. elegans* PIWI (PRG-1) to identify *in vivo* piRNA binding sites (9). Analysis of this data revealed that piRNAs bind both germline-silenced and germline-expressed mRNAs in *C. elegans*. Therefore, both bioinformatic and experimental approaches suggest that piRNAs alone are not sufficient to distinguish self from non-self nucleic acids. Importantly, CSR-1 Argonaute and its associated small RNAs have been shown to target germline-expressed RNAs and protect self nucleic acids from piRNA silencing (9,12,13). These studies suggest an interesting model where CSR-1 and its small RNAs act as a gene-licensing pathway, critical for distinguishing self nucleic acids from non-self in *C. elegans*.

Our genome-wide analyses of piRNA targeting sites have identified surprisingly few overlapping sites between experimentally-identified piRNA binding sites and sequence-based predicted piRNA targeting sites (10). These observations suggest that piRNA binding to mRNA targets may be regulated, such as by the CSR-1 pathway. However, how CSR-1 protects self RNAs from piRNA binding remains unknown. In addition, CSR-1 protection alone cannot fully explain gene licensing, as CSR-1 knockdown does not result in silencing of many CSR-1 targets (14,15) Currently it is unclear whether additional mechanisms exist to control piRNA recognition.

To gain insights into the mechanisms that regulate piRNA silencing, we examined the distribution of *in vivo* piRNA binding sites on target mRNAs. Surprisingly, we found that piRNAs preferentially bind to coding regions (CDS) of mRNAs. As a consequence, piRNAs trigger the production of secondary silencing small RNAs preferentially at the CDS. In contrast, our analyses showed that the sequence-based predicted piRNA targeting sites are slightly enriched in 3’ UTRs, thus the preference for CDS binding by piRNAs cannot be explained by the distribution of predicted piRNA targeting sites. In addition, we examined how anti-silencing Argonaute CSR-1 functions by analyzing piRNA binding sites from CSR-1 depleted animals. We found that CSR-1 inhibits both piRNA binding and secondary silencing small RNA production, but does so through distinct modes of action. Nonetheless, our analyses suggest CSR-1 is not responsible for restricting piRNA binding to the CDS. Together, our analyses identify CDSs as the critical regions that are uniquely susceptible to piRNA silencing in *C. elegans*.

## Results

### piRNAs exhibit a binding preference to the coding regions of target mRNAs in *C. elegans*

As described above, only a small proportion of predicted piRNA targeting sites were found to be bound by piRNAs *in vivo*, suggesting that piRNA binding to mRNA targets is regulated by cellular mechanisms (10). Intriguingly, while examining two well-characterized piRNA germline target mRNAs (9) (16), we noticed that despite the presence of several predicted piRNA targeting sites in various regions of these mRNAs, including CDS, 5’ UTRs and 3’ UTRs (Figure 1), the experimentally identified piRNA binding sites are mostly present in the CDSs, with some of these binding sites also predicted as piRNA targeting sites. We therefore wondered whether the observed piRNA CDS-binding preference could be a global trend, applicable to other germline mRNAs.

**Figure 1.**
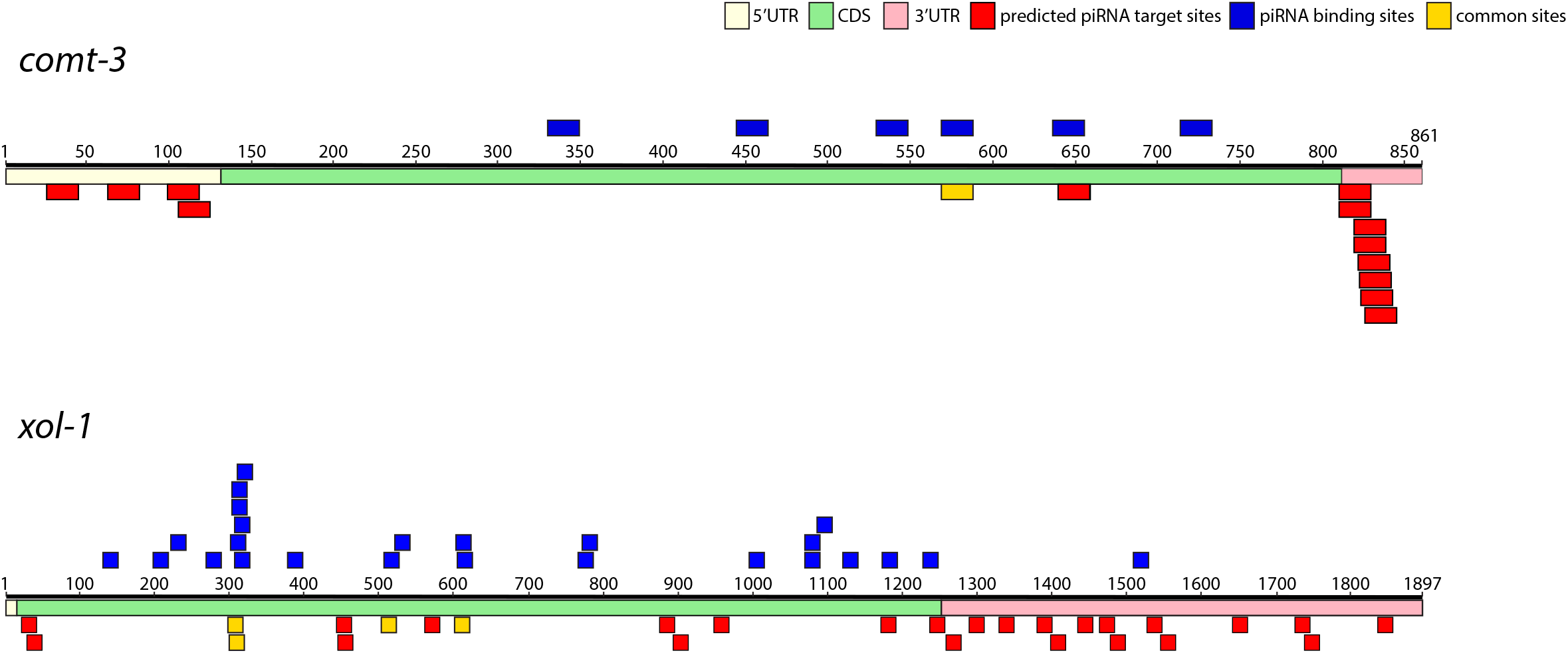
The distributions of predicted piRNA targeting sites and the experimentally identified piRNA binding sites on the indicated mRNAs. The location of predicted piRNA targeting sites are shown in red, the experimentally identified piRNA binding sites are shown in blue, and the common sites shared between predicted and experimentally identified sites are shown in yellow. The specific regions of the indicated mRNAs, including 5’ UTR, CDS and 3’ UTR are also labeled.

To test whether our analyses of Argonaute/small RNA binding sites are consistent with previously published results, we first examined the distribution of experimentally identified miRNA binding sites in *C. elegans*, which are known to be enriched in 3’ UTRs of mRNAs (17). Using previously published miRNA Argonaute ALG-1 iCLIP data, we identified transcriptome-wide *in vivo* miRNA binding sites and calculated the density of these binding sites in different regions of the mRNAs, including 5’ UTR, CDS or 3’ UTR (18). We observed a higher density for both the number of miRNA binding sites and the miRNA binding events at the 3’ end of mRNAs, especially at the 3’ UTR of mRNAs in *C. elegans* (Figure 2A & 2B and Figure S1A & S1B). To more closely examine the local change in miRNA binding preference between different regions, we examined the distribution of miRNA binding sites at the borders of CDS and UTRs. As expected, we observed that the number of miRNA binding sites did not change around the start codon but increased sharply immediately downstream of the stop codon (Figure 2C). Together, these analyses confirmed the previous observations that miRNAs predominately bind to their target mRNAs at their 3’ UTRs (17).

**Figure 2.**
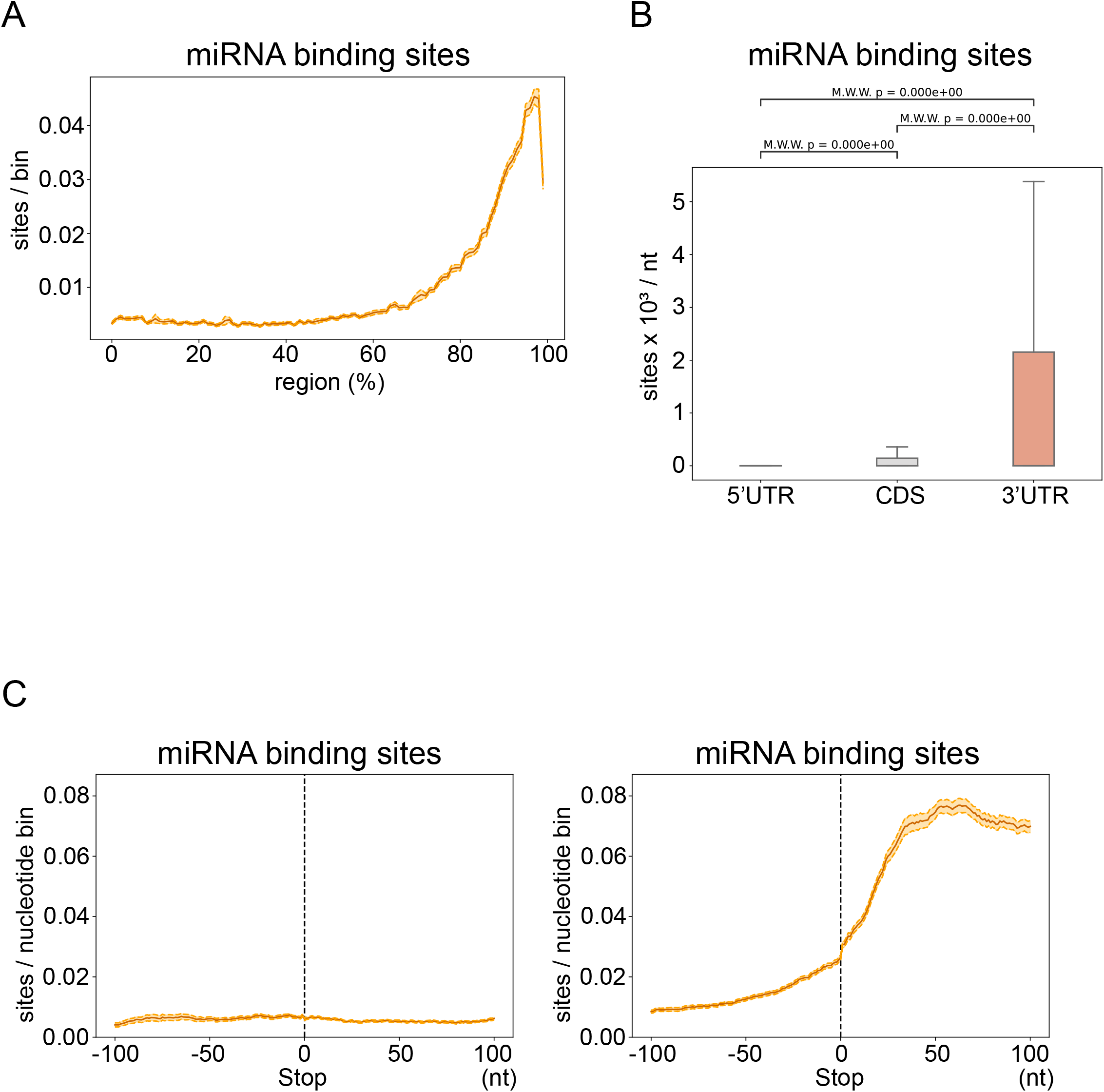
*In vivo* miRNA binding sites are enriched at the 3’ UTR of mRNAs in *C. elegans*. (A) Metagene trace shows the distribution of miRNA binding sites along an mRNA. The solid line indicates the average number of miRNA binding sites in all mRNAs. The dotted lines indicate the average number plus/minus one standard error. (B) The density of miRNA binding sites in the indicated regions. The statistical significance (P-value) of the difference in density between two regions was calculated by the Mann-Whitney U test. (C) The distribution of miRNA binding sites around the start codon (left) or stop codon (right). A 200 nucleotide-window centered at start or stop codon is shown. The solid line indicates the average number of miRNA binding sites in all mRNAs. For each position, the average is calculated from all mRNAs which possess that position. For example, not all mRNAs have 5’UTRs but they all have CDSs. The dotted lines indicate the average number plus/minus one standard error.

We then compared the genome-wide distribution of sequence-based predicted piRNA targeting sites and experimentally identified piRNA binding sites on germline mRNAs, since piRNAs are mostly expressed in the germline (19,20). Using the stringent piRNA targeting rule identified in previous studies (8,19), we observed that the predicted piRNA targeting sites are enriched at the 3’ end of germline mRNAs (Figure 3A), and their densities are highest in the 3’ UTR of germline mRNAs (Figure 3B). Using published PIWI PRG-1 CLASH data (9), we then analyzed the transcriptome-wide *in vivo* piRNA binding sites on germline mRNAs (18). Interestingly, we observed a very distinct distribution of *in vivo* piRNA binding sites compared to that of the predicted piRNA targeting sites; the overall distribution of piRNA binding sites on germline mRNAs exhibits a peak at the 5’ end, followed by a steady level across the gene body before a decrease at the 3’ end (Figure 3C). Consistent with the CDS binding preference found in the well-characterized piRNA targeted mRNAs, we found that the density of *in vivo* piRNA binding sites is highest in the CDS of germline mRNAs (Figure 3D). Specifically, the piRNA binding sites increased most notably around the start codon and decreased most notably around the stop codon (Figure 3E) – opposite to the distribution of the predicted piRNA targeting sites (Figure S2A). These observations suggest that piRNAs preferentially bind to CDSs, and this preference cannot be explained by the distribution of predicted piRNA targeting sites.

**Figure 3.**
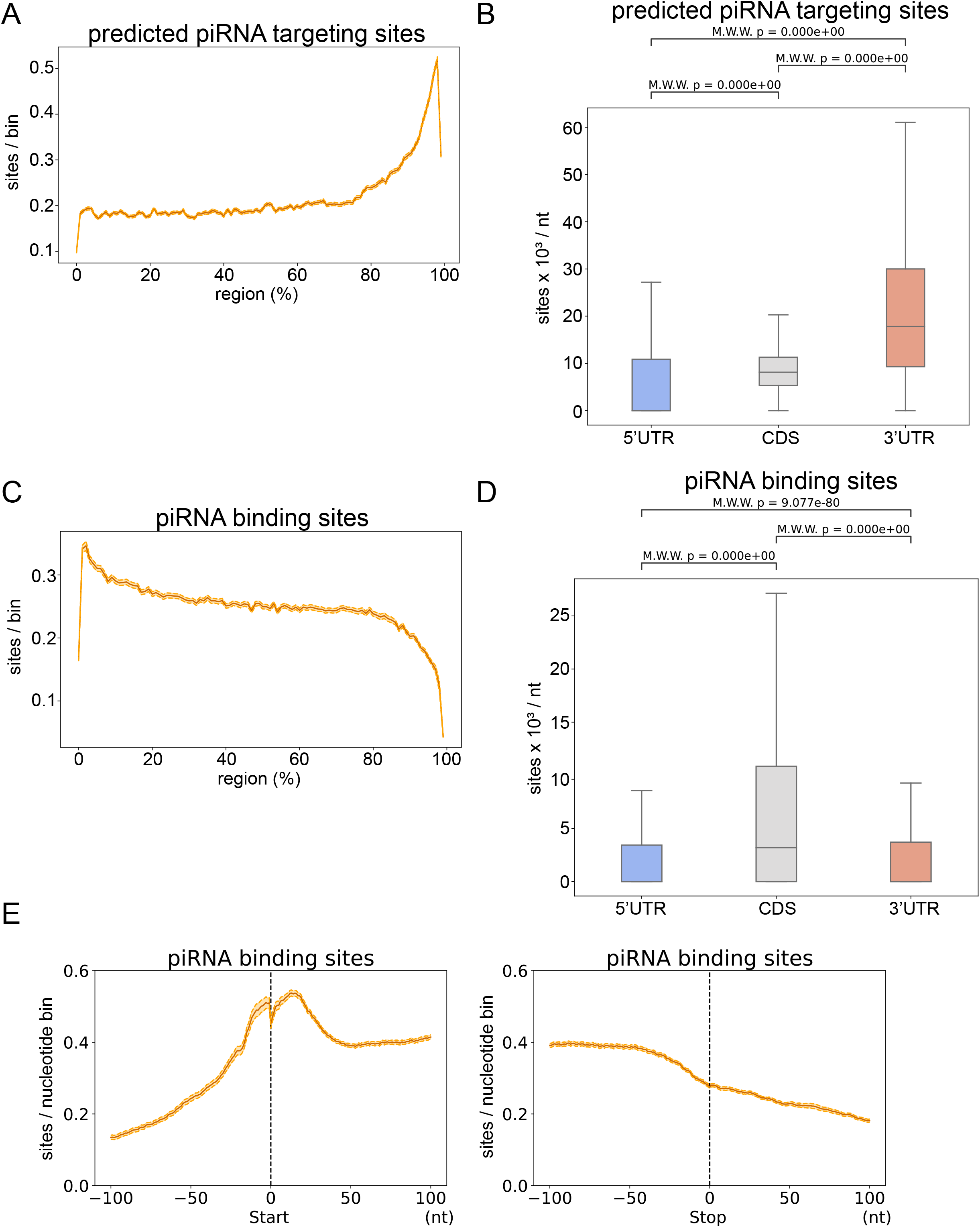
*In vivo* piRNA binding sites are enriched in the coding region (CDS) of germline mRNAs in *C. elegans*. (A) Metagene trace shows the distribution of predicted piRNA targeting sites along a germline mRNA. The solid line indicates the average number of piRNA targeting sites in all germline mRNAs. The dotted lines indicate the average number plus/minus one standard error. (B) The density of predicted piRNA targeting sites in the indicated regions of germline mRNAs. The statistical significance (P-value) of the difference in density between two regions was calculated by the Mann-Whitney U test. (C) Metagene trace shows the distribution of experimentally identified piRNA binding sites along a germline mRNA. The solid line indicates the average number of piRNA binding sites in all germline mRNAs. The dotted lines indicate the average number plus/minus one standard error. (D) The density of experimentally identified piRNA binding sites in the indicated regions of germline mRNAs. The statistical significance (P-value) of the difference in density between two regions was calculated by the Mann-Whitney U test. (E) The distribution of piRNA binding sites around the start codon (left) or stop codon (right). A 200 nucleotide-window centered at start or stop codon is shown. The solid line indicates the average number of piRNA binding sites in all germline mRNAs. For each position, the average is calculated from all germline mRNAs which possess that position.

As some miRNAs are also expressed in the germline, we wondered whether the enrichment of piRNA binding sites in CDSs is a feature shared by germline-expressed miRNAs (21). We therefore restricted our miRNA binding site analysis to germline-enriched miRNAs at germline mRNAs. However, we found that miRNA binding sites for these germline-enriched miRNAs remain enriched at the 3’ UTR and 3’ end of germline mRNAs (Figure S2B). Therefore, the piRNA CDS-binding preference is not shared by miRNAs in the germline. Since previous studies have shown that germline-expressed transcripts are protected from piRNA silencing by CSR-1 and its associated small RNAs (9,12,13), we also examined whether there is a difference in piRNA binding distribution on germline-expressed mRNAs (here defined as CSR-1 targets) or germline-silenced mRNAs (here defined as WAGO targets, see more descriptions of WAGO Argonautes below). We found that the CDS enrichment for piRNA binding sites exists for both WAGO targets as well as for CSR-1 targets (Figure S2C) (22,23). This observation suggests that the CDS binding preference is likely not caused by CSR-1 (see more about CSR-1 function below). Together, we conclude that the *in vivo* piRNA binding sites are enriched in the CDSs of germline-expressed mRNAs in *C. elegans*.

### piRNAs preferentially trigger the production of secondary small RNAs at mRNA coding regions

piRNAs induce gene silencing by triggering the production of secondary silencing small RNAs that associate with Worm specific ArGOnautes (WAGOs), also known as WAGO 22G-RNAs (24–26). These WAGO 22G-RNAs are produced by RNA-dependent RNA polymerase locally at piRNA binding sites. Since piRNAs preferentially bind to CDSs, we wondered whether the production of WAGO 22G-RNAs is also enriched at CDS regions. To test this hypothesis, we compared the density of WAGO 22G-RNAs in distinct regions of germline mRNAs. Indeed, for germline-silenced mRNAs (WAGO targets) (22), we observed that both WAGO-1 and WAGO-9 (also known as HRDE-1) 22G-RNAs are significantly enriched in the CDS (Figure 4A and data not shown). In addition, for germline-expressed mRNAs (CSR-1 targets), WAGO 22G-RNAs are also enriched in the CDS (Figure 4A), despite their overall WAGO 22G-RNA levels being much lower in CSR-1 targets than in WAGO targets. WAGO 22G-RNA production can be induced by piRNAs and other small RNA pathways (27,28). Therefore, if piRNA-induced WAGO 22G-RNAs are produced preferentially at the CDS, we expect that in PIWI mutants, which lose all piRNAs, the WAGO 22G-RNAs should be preferentially reduced at coding regions. Indeed, when we compared the levels of WAGO-1 22G-RNAs between wild type and PIWI *prg-1* mutants (29), we observed a greater reduction of WAGO-1 22G-RNAs in the CDS than that in 5’ or 3’ UTRs, for both germline-silenced and germline-expressed mRNAs (Figure 4B). These observations suggest that the preferential piRNA binding at the CDS of mRNAs is functionally relevant, as piRNA binding preference in the CDS leads to a corresponding enrichment of secondary WAGO 22G-RNAs produced at the CDS. Interestingly, a recent study compared the ability of synthetic piRNAs to trigger gene silencing and showed that those synthetic piRNAs targeting the CDS can trigger robust gene silencing, but for synthetic piRNA targeting 5’ or 3’UTRs, gene silencing is not a successful (30). These reporter-based experiments support our transcriptome-wide analyses that the CDS is the critical region that is uniquely competent for piRNA silencing in *C. elegans*.

**Figure 4.**
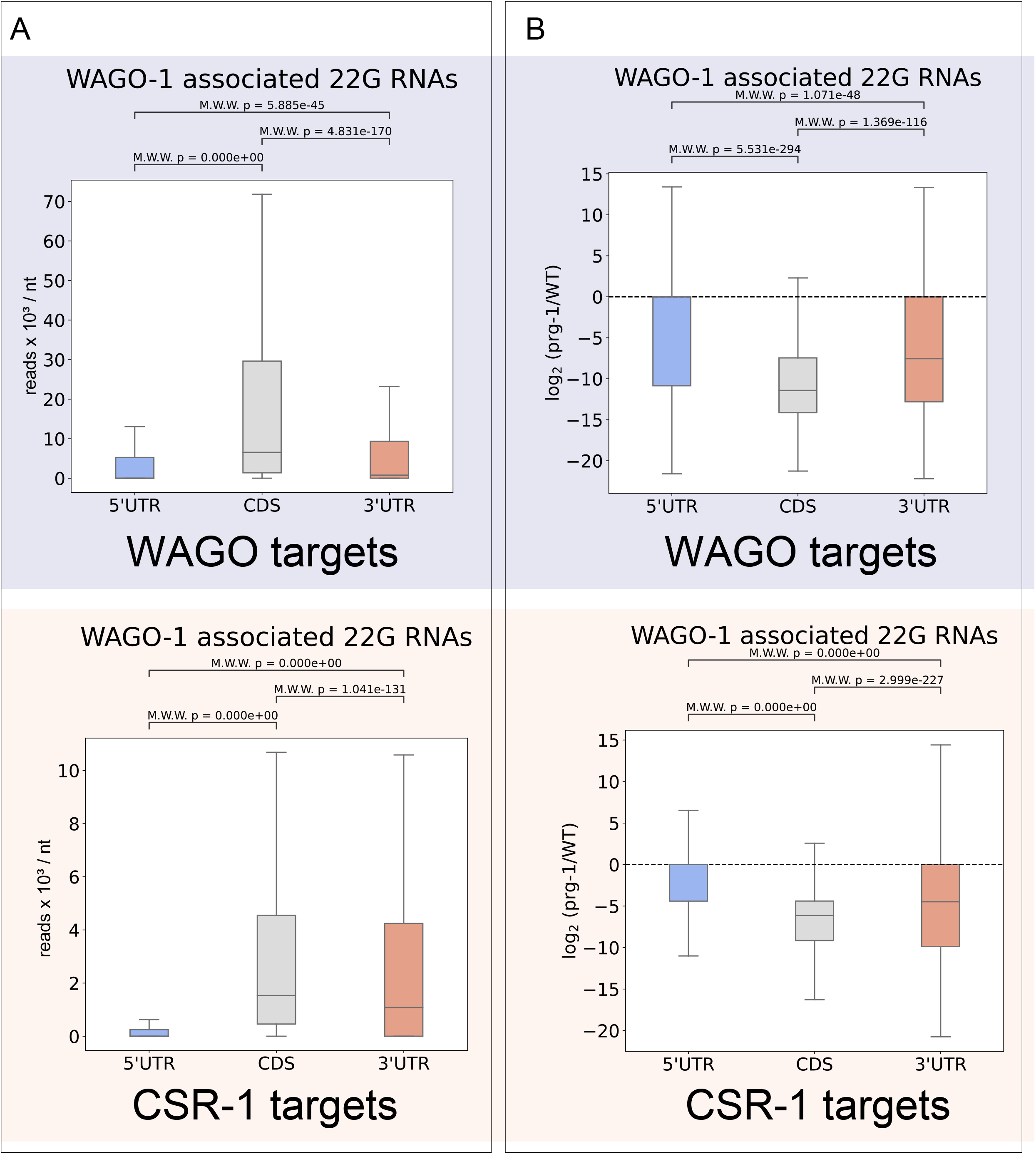
Secondary WAGO 22G-RNAs are preferentially produced in coding regions. (A) The density of WAGO-1 22G-RNAs in the indicated regions of WAGO targets (top) or CSR-1 (bottom) targets. The statistical significance (P-value) of the difference in density between two regions was calculated by the Mann-Whitney U test. (B) The ratio of WAGO-1 22G-RNAs in the *prg*-1 mutant over those in wild type in the indicated regions of WAGO (top) or CSR-1 (bottom) targets. The statistical significance (P-value) of the difference in fold change (*prg-1* mutant/WT) between two regions was calculated by the Mann-Whitney U test.

### Distinct modes of action for CSR-1 to inhibit piRNA binding and WAGO 22G-RNA production

CSR-1 Argonaute and its associated 22G-RNAs have been reported to protect germline transcripts from piRNA silencing (12,13). However, the mechanism behind CSR-1 Argonaute’s anti-piRNA silencing function remains unclear. A previous study suggests that CSR-1 inhibits the piRNA pathway by interfering with piRNA binding (9). Nonetheless, it is unclear whether CSR-1 also inhibits downstream WAGO 22G-RNA synthesis. In addition, it is unknown whether CSR-1 binding leads to protection of the whole transcript from piRNA silencing, or if CSR-1 only interferes with piRNA silencing locally, proximal to CSR-1 targeting sites. To gain insights into CSR-1’s anti-piRNA silencing function, we first examined the distribution of CSR-1 22G-RNAs on germline mRNAs. We found that the levels of CSR-1 22G-RNAs are higher at the 3’ end of CSR-1 targets (Figure 5A, top) and are produced at both CDSs and 3’ UTRs (Figure S3A, top), consistent with a recent report (15). In addition, we found a low level of CSR-1 22G-RNAs produced from WAGO targets. These CSR-1 22G-RNAs are more equally distributed across the gene body, with their levels reduced at both 5’ and 3’ ends (Figure 5A, bottom), and enriched at the CDS (Figure S3A, bottom).

**Figure 5.**
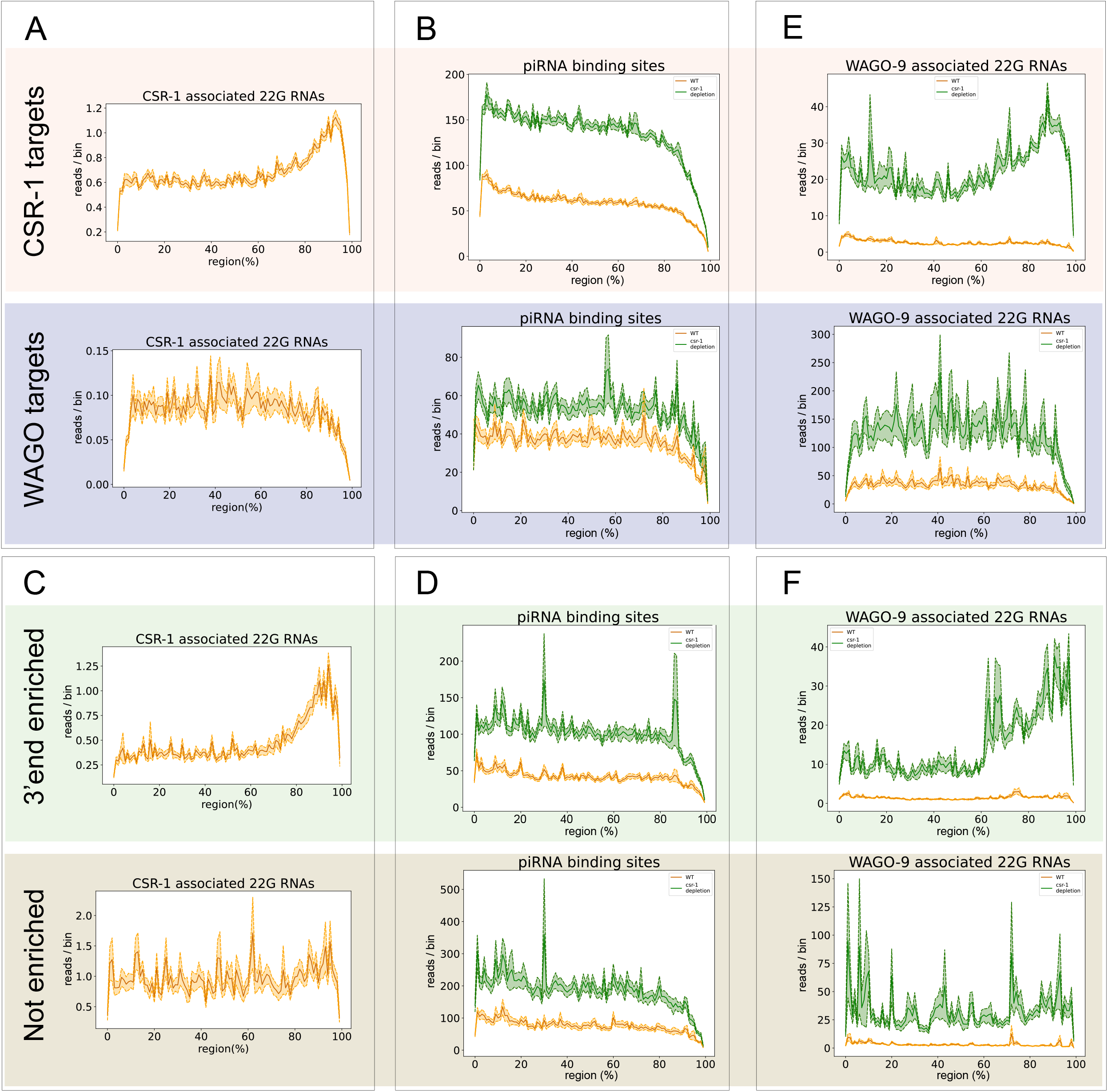
CSR-1 inhibits piRNA binding and WAGO 22G-RNA synthesis with distinct modes of action. (A) Metagene trace shows the distribution of CSR-1 22G-RNAs along CSR-1 (top) or WAGO (bottom) targets. The solid line indicates the average number of CSR-1 22G-RNAs read counts in germline mRNAs. The dotted lines indicate the average number plus/minus one standard error. (B) Metagene trace shows the distribution of piRNA binding events in wild type (orange) or in CSR-1 depleted animals (green) mapped to CSR-1 (top) or WAGO (bottom) targets. The solid line indicates the average number of piRNA binding events (read counts) in CSR-1 or WAGO targets. The dotted lines indicate the average number plus/minus one standard error. (C) Metagene trace shows the distribution of CSR-1 22G-RNAs mapped to the indicated CSR-1 subgroup genes, including 3’ end 22G enriched (top) or not enriched (bottom) CSR-1 targets. The solid line indicates the average number of CSR-1 22G-RNAs read counts in the indicated CSR-1 subgroup genes. The dotted lines indicate the average number plus/minus one standard error. (D) Metagene trace shows the distribution of piRNA binding events in wild type (orange) or in CSR-1 depleted animals (green) mapped to the indicated CSR-1 subgroup genes, including 3’ end 22G enriched (top) or not enriched (bottom) CSR-1 targets. The solid line indicates the average number of piRNA binding events (read counts) in the indicated CSR-1 subgroup genes. The dotted lines indicate the average number plus/minus one standard error. (E) Metagene traces show the distribution of WAGO-9 (HRDE-1) 22G-RNAs in wild type (orange) or in CSR-1 depleted animals (green) mapped to CSR-1 (top) or WAGO (bottom) targets. The solid line indicates the average number of WAGO-9 22G-RNA read counts in CSR-1 or WAGO targets. The dotted lines indicate the average number plus/minus one standard error. (F) Metagene traces show the distribution of WAGO-9 (HRDE-1) 22G-RNAs in wild type (orange) or in CSR-1 depleted animals (green) mapped to the indicated CSR-1 subgroup genes, including 3’ end 22G enriched (top) or not enriched (bottom) CSR-1 targets. The solid line indicates the average number of WAGO-9 22G-RNAs read counts in the indicated CSR-1 subgroup genes, including 3’ end 22G enriched (top) or not enriched (bottom) CSR-1 targets. The dotted lines indicate the average number plus/minus one standard error.

To examine the relationship between CSR-1 binding and piRNA binding, we then examined the effect of CSR-1 depletion on piRNA binding (9). Our analyses showed that CSR-1 depletion leads to a greater increase of piRNA binding in CSR-1 targets than in WAGO-1 targets, indicating that the greater level of CSR-1 22G-RNA production leads to greater levels of protection from piRNA binding (Figure 5B and Figure S3B). If CSR-1 inhibits piRNA binding locally, we expect a greater increase of piRNA binding at regions where CSR-1 levels are higher. However, despite the very different distribution of CSR-1 22G-RNA accumulation on WAGO targets and CSR-1 targets, we observed a similar increase of piRNA binding in all regions of the transcripts for both WAGO and CSR-1 targets (Figure 5B and Figure S3B) upon CSR-1 depletion. Therefore, while the overall level of CSR-1 targeting a given transcript correlates with its protection level against piRNA binding, the location of CSR-1 targeting within the transcript does not seem to matter. To further examine the relationship between CSR-1 localization and piRNA binding, we defined subgroups of CSR-1 targets for CSR-1 targets where CSR-1 22G-RNAs are either enriched at their 3’ UTRs (n=2082) or are not enriched at any specific regions (n=763) (Figure 5C and Figure S3C). Again, upon CSR-1 depletion, we found that piRNA binding sites increased in all regions of both subgroups of CSR-1 targets despite their different CSR-1 distributions (Figure 5D and Figure S3D). These results suggest that CSR-1’s inhibition of piRNA binding does not correlate with CSR-1 22G-RNA distribution.

Surprisingly, we observed that piRNAs bind germline-expressed CSR-1 targets with higher density than they bind germline-silenced WAGO targets (Figure S3E). Significantly more WAGO 22G-RNAs (both WAGO-1 and WAGO-9 22G-RNAs) were produced from all regions of WAGO targets compared to CSR-1 targets (Figure S3F), indicating that piRNA binding on CSR-1 targets does not trigger WAGO 22G-RNA production as effectively as piRNA binding on WAGO targets. This observation suggests that CSR-1 not only inhibits piRNA binding, it can also inhibit the downstream production of WAGO 22G-RNAs. To examine the relationship between CSR-1 binding and WAGO 22G-RNA synthesis, we examined the change of WAGO-9 (HRDE-1) 22G-RNAs upon CSR-1 depletion (15). Interestingly, for both CSR-1 and WAGO targets, there is a greater increase in WAGO-9 22G-RNAs preferentially at regions where CSR-1 22G RNAs are enriched, including the 3’ end of CSR-1 targets and at the gene body of WAGO targets, respectively (Figure 5E and Figure S3G). To further examine the relationship between CSR-1 distribution and WAGO 22G-RNA production, we compared those CSR-1 target subgroups mentioned above that have either CSR-1 22G-RNAs enriched at 3’ UTRs or not enriched at a specific region. Upon CSR-1 depletion, we found a greater increase in WAGO 22G-RNA production at 3’ UTRs only for those CSR-1 targets with a 3’ CSR-1 targeting enrichment (Figure 5F and Figure S3H). These observations show that the local CSR-1 22G-RNA levels contribute to the inhibitory strength against WAGO 22G-RNA production, indicating that CSR-1 acts locally to inhibit the production WAGO 22G-RNAs. Together, our analyses suggest that CSR-1 can counteract the piRNA pathway through the inhibition of both piRNA binding and WAGO 22G-RNA production. In addition, our analyses support a model where CSR-1 provides transcript-wide protection against piRNA binding, but only acts locally to inhibit WAGO 22G-RNA production (Figure 6).

**Figure 6.**
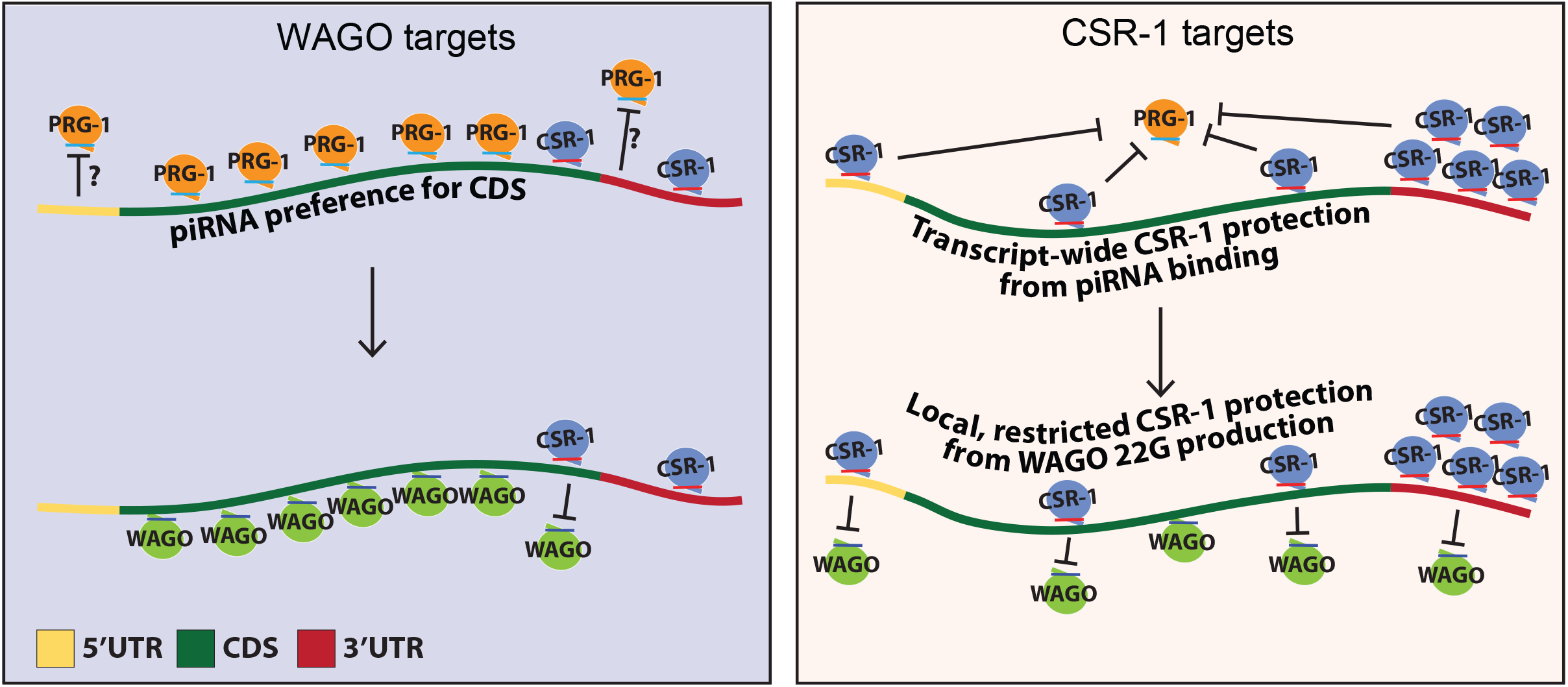
A model depicting piRNAs’ CDS binding preference and 22G-RNA production on WAGO and CSR-1 targets. For WAGO targets (left), piRNAs exhibit a preference for targeting the CDS which is independent of CSR-1. WAGO production which is initiated by piRNA targeting follows a similar pattern of enrichment. For CSR-1 targets (right), CSR-1 blocks piRNA targeting in a transcript-wide mode of action. For downstream WAGO production, CSR-1 acts locally to block WAGO accumulation.

### CSR-1 is not responsible for piRNAs’ CDS-binding preference

We then investigated whether CSR-1 contributes to the piRNA CDS binding preference. If CSR-1 is responsible for the piRNA CDS-binding preference, we would expect this preference to diminish upon CSR-1 depletion. However, upon CSR-1 knockdown, piRNAs retain their CDS binding preference for both CSR-1 targets and WAGO targets (Figure S3B). Specifically, the knockdown of CSR-1 did not change the pattern of piRNA binding sites around start or stop codons for either CSR-1 targets or WAGO-1 targets (Figure S4A). As translation has shown to antagonize CSR-1 22G-RNA synthesis in CDS regions (15), we wondered whether translation may also play a role in regulating piRNA binding to the CDS. For both highly-translated and lowly-translated CSR-1 or WAGO-1 targets (31), we found that piRNAs retain their CDS binding preference (Figure S4B). However, since translation of germline mRNAs is a highly regulated process throughout germline development, future studies are needed to further investigate whether translation plays a role in regulating piRNA binding. Together, our analyses suggest that the piRNA CDS binding preference cannot be explained by CSR-1 licensing and indicates that an unknown mechanism promotes or restricts piRNA binding to the CDS.

## Discussion

Small RNAs, such as miRNAs and piRNAs, regulate gene expression through base-pairing with their target mRNAs. It has been shown that miRNA targeting sites are enriched at 3’ UTRs and that 3’ UTR sites exhibit more gene regulatory effects than those at the CDS (32). While piRNAs are known for their roles in genome defense, it was previously unknown whether piRNA targeting or regulatory potential operate differently among the different regions of their mRNA targets. By analyzing transcriptome-wide *in vivo* piRNA binding sites, we uncovered a new feature regulating the piRNA defense system in *C. elegans: C. elegans* piRNAs preferentially bind and trigger gene silencing at the CDS of germline mRNAs. Consequently, secondary WAGO 22G-RNAs are preferentially produced at the CDS as well (Figure 6). Our finding has important implications on using synthetic piRNAs to silence target genes of interest, where piRNAs targeting the CDS are likely going to be more effective. Indeed, a recent publication has confirmed this trend, albeit not on a transcriptome-wide scale (30). Nonetheless, it is worth noting that synthetic piRNAs targeting the 3’ UTRs of transgenes can trigger transgene silencing (25,26). In addition, the CDS of *oma-1*, but not its UTRs has been reported to carry specific licensing properties that counter piRNA silencing (33). Future studies will be necessary to further understand the underlying mechanisms that confer such differences in sensitivity to piRNA silencing. Furthermore, our study also provides insights into the applications in the expression of foreign nucleic acids in the germline of *C. elegans*. piRNAs are known to trigger gene silencing of transgenes that carry foreign nucleic acids, and removal of piRNA sites from the coding regions of these foreign nucleic acids are thus likely to be critical for the successful expression of these foreign nucleic acids in *C. elegans*. Indeed, previous studies have optimized the coding regions of foreign nucleic acids, including GFP, mCherry or Cas9, which successfully prevent these silencing-prone transgenes from piRNA-induced gene silencing (8).

Our work also provides insights into the mechanisms by which CSR-1 inhibits the piRNA pathway. Specifically, we found that CSR-1 depletion leads to increased piRNA binding across whole CSR-1 targeting transcripts, while increased secondary WAGO 22G-RNA synthesis preferentially occurs locally at CSR-1 targeting sites. These observations suggest that the mechanism underlying these two CSR-1 anti-silencing processes is different. Our analyses indicate that CSR-1 binding at various locations in a transcript can result in the protection of the entire transcript from piRNA recognition (Figure 6). Indeed, a recent report has shown that tethering of CSR-1 to a single locus at the 3’ UTR of an endogenous mRNA can protect that mRNA from piRNA silencing (34). In addition, our analyses reveal that CSR-1 targeted regions are locally protected against WAGO 22G-RNA production (Figure 6). Interestingly, previous analyses on transgenes have indicated a local inhibition of WAGO 22G-RNAs in CSR-1 targeted regions; in transgenic worms carrying a silenced GFP::CDK-1 transgene, only foreign GFP sequences produce high levels of WAGO 22G-RNA, while the CDK-1 sequence (which is identical to the endogenous CDK-1 gene and therefore targeted by CSR-1 22G-RNAs) produces low levels of WAGO 22G-RNAs, indicating a local inhibition of WAGO 22G-RNAs at the CSR-1 targeted region (13). One interesting working model is that CSR-1 binding can somehow remove its bound mRNA from loci of piRNA recognition, while CSR-1 inhibits WAGO 22G-RNAs synthesis locally, such as by competing for RNA-dependent RNA polymerases – resources known to be shared by these two Argonautes to produce their associated small RNAs (22).

Finally, our analyses also suggest that the CSR-1 pathway is not responsible for piRNAs’ CDS binding preference, suggesting that an unknown mechanism promotes CDS surveillance by piRNAs. As the CDSs of foreign RNAs are expected to be under more selective pressure than UTRs, we speculate that the preferential recognition of CDSs by piRNAs likely offers a mechanism for the piRNA surveillance system to reduce the chance of evolution of foreign RNA variants that evade piRNA recognition.

## Conclusions

We analyzed transcriptome-wide *in* vivo piRNA binding sites and found that *C. elegans* piRNAs preferentially bind the CDS of germline mRNAs. We found that CDS binding events lead to a significant enrichment in siRNA production templating CDSs. Surprisingly, licensing Argonaute CSR-1 does not drive the piRNA binding enrichment, but CSR-1 interaction does protect transcripts globally from PRG-1 binding and locally from WAGO targeting. Our work will have important ramifications for targeted gene silencing in *C. elegans*, and it may reveal strategies evolved in the piRNA pathway to continue to specifically silence its targets in spite of genetic drift.

## Methods

### Analyses of in vivo miRNA and piRNA binding sites

The iCLIP data of ALG-1 (SRR3882949) and the CLASH data of PRG-1 (SRR6512652/WT and SRR6512654/CSR-1 depletion) were used in these analyses (9,17). To identify *in vivo* miRNA and piRNA binding sites, hybrids of miRNA/piRNA with their target mRNAs were identified by CLASH analyst (18) with default settings for data pre-processing and searching algorithms. The *C. elegans* miRNA/piRNA sequences and the mRNA sequences (n=43040) of Wormbase version WS275 were used. Once hybrid reads are identified, miRNA and piRNA binding sites are defined by the regions of interacting mRNAs with lowest binding energy or highest binding score to corresponding miRNA using RNAup (35) or to corresponding piRNA using pirScan (19), respectively. When the mRNA interacting sequences (CLASH identified regions) are shorter than miRNA or piRNA, they are first extended to the size of miRNA and piRNAs using both the upstream and downstream sequences before they are examined for sites with best pairing energy/score.

Germline mRNAs (n=27792) are defined as mRNAs that are detected either in the spermatogenic or oogenetic transcriptome (20). WAGO targets (n=3644) are defined as transcripts whose mapped 22G-RNAs exhibit over two-fold enrichment from either WAGO-1 IP than that from input 22G-RNAs (22) or WAGO-9 IP than that from input 22G-RNAs (36). CSR-1 targets (n=15821) are defined as transcripts whose mapped 22G-RNAs exhibit over two-fold enrichment from CSR-1 IP than that from input 22G-RNAs (23).

### Prediction of piRNA targeting sites

Predicted piRNA targeting sites (with stringent targeting rules) on *C. elegans* germline mRNAs were obtained from piRTarbase (10). For stringent rules, the following criteria are used; at seed region, no non-GU mismatches and no more than 2 GU mismatches are allowed. At non-seed region, no more than 2 non-GU mismatches and no more than 3 GU mismatches are allowed. Furthermore, no more than 6 total mismatches are allowed from both seed and non-seed regions (19).

### Measurements of miRNA, piRNA binding sites and 22G-RNAs levels at different regions of mRNAs

The *C. elegans* transcriptome data (WS275) annotation was used to define the location of 5’ UTRs, CDSs, and 3’ UTRs. The number of sites or read counts from each region were then divided by the nucleotide length of the corresponding regions for each mRNA to obtain the density. Where sites were spanning between two distinct regions, the sites or read counts were split according to the portion that mapped to each region. For measurements of sites/reads around start and stop codons, the 200 nt (+/- 100 nt) window centered at start or stop codon of each transcript was aligned and the number of the sites/reads per transcript mapped at each nucleotide was calculated. As transcripts have different 5’ and 3’UTR sizes, the mapped sites/read counts at each position were divided by the number of transcripts that contain such position to obtain the sites/read counts per transcript. The following small RNA data were used to analyze the distribution and levels of CSR-1 and WAGO 22G-RNAs; CSR-1 associated small RNAs data (SRR12318132), WAGO-1 associated small RNAs (SRR8482951/WT, SRR8482949/*prg-1* mutant) (29), and WAGO-9 (HRDE-1) associated small RNAs (SRR12318140/WT, SRR12318144/CSR-1 depletion) (15).

### Metagene analyses

A custom script was used to divide mRNA transcripts into 100 bins and the normalized reads within each bin was calculated. The average number of sites or read counts at each bin per mRNA transcript was then calculated.

### Ribo-seq analyses

The ribo-seq data from the wild type *C. elegans* late L4/young adult sample (36 hours post hatching, SRR3356499) (31) were used to identify the number of ribosome protected fragments (RPF) mapped to each CSR-1 target or WAGO-1 target. The transcripts with top 10% or bottom 10% amounts of RPF were defined as highly-translated and poorly-translated transcripts, respectively.

## Acknowledgements

This work is supported in part by NIH predoctoral training grant T32 GM07197 to J.B.; the Ministry of Science of Technology of Taiwan (MOST 108-2628-E-006-004-MY3 and MOST 110-2221-E-006-198-MY3 grants to W.-S.W., the NIH grant R01-GM132457 to H.-C.L.

## Data availability statement

All sequencing data analyzed in the manuscript are available at NCBI GEO or ENA database. The SRR numbers of sequencing data used in specific analyses are provided in the Materials and Methods section above.

