## Supplemental information for "Transcriptome-wide analysis suggests piRNAs preferentially recognize the coding region of mRNAs in *C. elegans*"

**This supplementary information includes:**

**Figure S1.** ***In vivo* miRNA binding sites are enriched at the 3' UTR of mRNAs in *C. elegans*, related to Figure 2.**

**Figure S2. piRNAs' CDS binding preference cannot be explain by the predicted piRNA targeting sites, related to Figure 3.**

**Figure S3. The distribution of CSR-1 22G-RNAs, WAGO 22G-RNAs, and piRNA binding sites, related to Figure 5.**

**Figure S4. The distribution of piRNA binding sites, related to Figure 5.**

**Figure S1.** ***In vivo*** **miRNA binding sites are enriched at the 3' UTR of mRNAs in** ***C. elegans*****, related to Figure 2.**

(A) Metagene trace shows the distribution of miRNA binding events along an mRNA. The solid line indicates the average number of miRNA binding events (read counts) in all mRNAs. The dotted lines indicate the average number plus/minus one standard error.

(B) The density of miRNA binding events in the indicated regions. The statistical significance (P-value) of the difference in density between two regions was calculated by the Mann-Whitney U test.

**Figure S2. piRNAs' CDS binding preference cannot be explain by the predicted piRNA targeting sites, related to Figure 3.**

(A) The distribution of predicted piRNA targeting sites around the start codon (left) or stop codon (right). A 200 nucleotide-window centered at start or stop codon is shown. The solid line indicates the average number of predicted piRNA targeting sites in all germline mRNAs. For each position, the average is calculated from all germline mRNAs which possess that position.

(B) Metagene trace shows the distribution of germline-expressed miRNA binding sites along a germline mRNA (left). The solid line indicates the average number of miRNA binding events (read counts) in germline mRNAs. The dotted lines indicate the average number plus/minus one standard error. The density of experimentally identified germline-expressed miRNA binding sites in the indicated regions of germline mRNAs (right). The statistical significance (P-value) of the difference in density between two regions was calculated by the Mann-Whitney U test.

(C) The density of piRNA binding sites in the indicated regions of germline-silenced mRNAs (WAGO targets, left) and germline-expressed mRNAs (CSR-1 targets, right). The statistical significance (P-value) of the difference in density between two regions was calculated by the Mann-Whitney U test.

**Figure S3. The distribution of CSR-1 22G-RNAs, WAGO 22G-RNAs, and piRNA binding sites, related to Figure 5**

(A) The density of CSR-1 22G-RNAs in the indicated regions of CSR-1 (top) or WAGO (bottom) targets. The statistical significance (P-value) of the difference in density between two regions was calculated by the Mann-Whitney U test.

(B) The density of piRNA binding events in the indicated regions in wild type (orange) or in CSR-1 depleted animals (green) mapped to CSR-1 (top) or WAGO (bottom) targets. The statistical significance (P-value) of the difference in density between two regions was calculated by the Mann-Whitney U test.

(C) The density of CSR-1 22G-RNAs mapped to the indicated regions of CSR-1 subgroup genes, including 3’ end 22G-RNA enriched (top) or not enriched (bottom) CSR-1 genes. The statistical significance (P-value) of the difference in density between two regions was calculated by the Mann-Whitney U test.

(D) The density of piRNA binding events mapped to the indicated regions in wild type (orange) or in CSR-1 depleted animals (green) of CSR-1 subgroup genes, including 3’ end 22G-RNA enriched (top) or not enriched (bottom) CSR-1 genes. The statistical significance (P-value) of the difference in density between two regions was calculated by the Mann-Whitney U test.

(E) The density of piRNA binding events (read counts) in the indicated regions of WAGO targets or CSR-1 targets. The statistical significance (P-value) of the difference in density between two regions was calculated by the Mann-Whitney U test.

(F) The density of WAGO-9 (left) or WAGO-1 (right) 22G-RNAs mapped to the regions of WAGO targets or CSR-1 targets. The statistical significance (P-value) of the difference in density between two regions was calculated by the Mann-Whitney U test.

(G) The density of WAGO-9 (HRDE-1) 22G-RNAs mapped to the indicated regions in wild type or in CSR-1 depleted animals of CSR-1 (top) or WAGO targets. The statistical significance (P-value) of the difference in density between two regions was calculated by the Mann-Whitney U test.

(H) The density of WAGO-9 (HRDE-1) 22G-RNAs in wild type or in CSR-1 depleted animals mapped to the indicated regions of CSR-1 subgroup genes, including 3’ end 22G-RNA enriched (top) or not enriched (bottom) CSR-1 genes. The statistical significance (P-value) of the difference in density between two regions was calculated by the Mann-Whitney U test.

**Figure S4. The distribution of piRNA binding sites, related to Figure 5**

(A) The distribution of piRNA binding events around the start codon (left) or stop codon (right) in wild type or in CSR-1 depleted animals. A 200 nucleotide-window centered at start or stop codon is shown. The solid line indicates the average number of WAGO-9 22G-RNAs read counts in CSR-1 targets or WAGO-1 targets.

(B) The density of piRNA binding events (read counts) in the indicated regions of highly-translated or poorly-translated WAGO targets (left) and highly-translated or poorly-translated CSR-1 targets (right). The statistical significance (P-value) of the difference in density between two regions was calculated by the Mann-Whitney U test.

### Figure S1

A

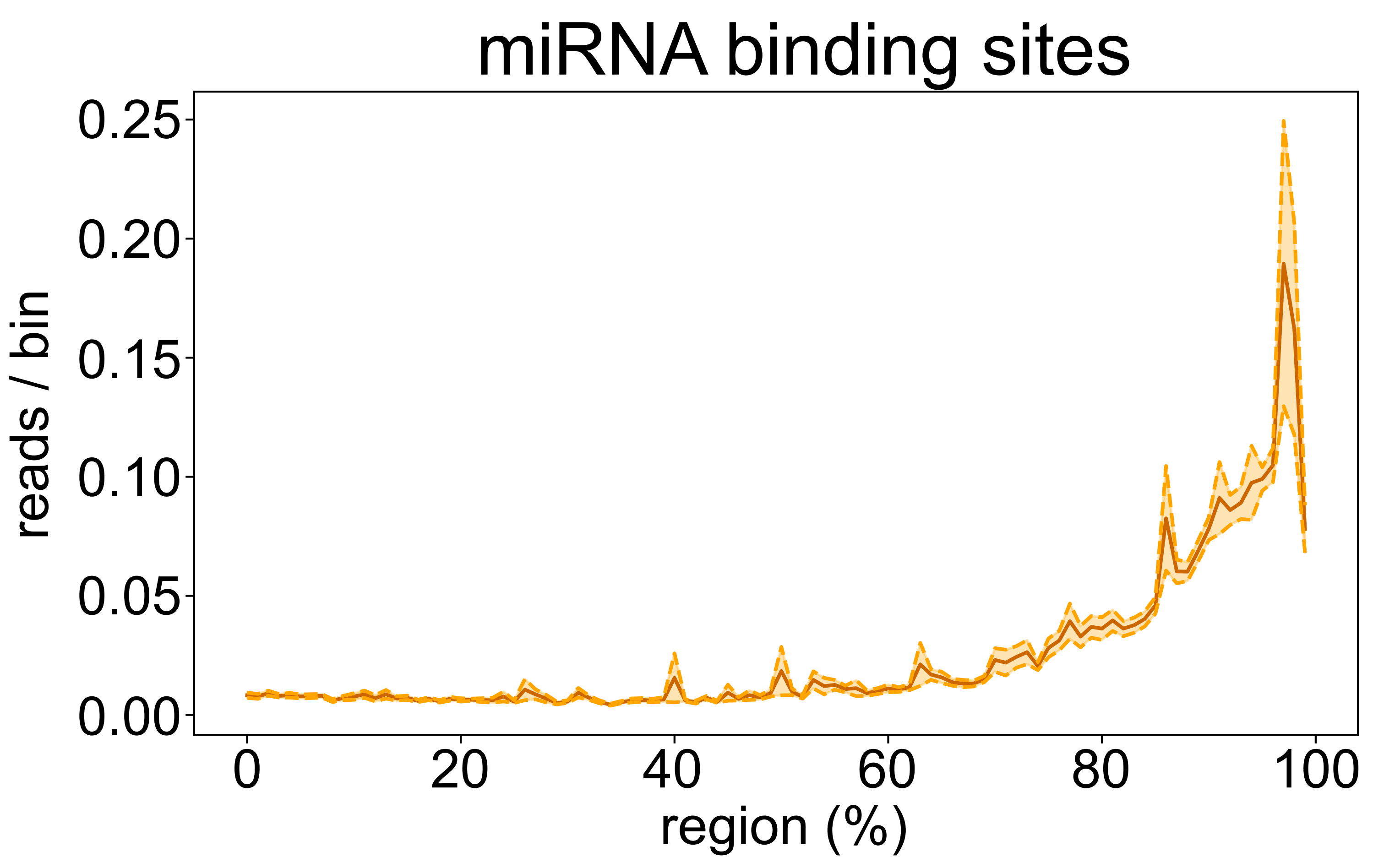
miRNA binding sites
reads / bin
0.00
0.05
0.10
0.15
0.20
0.25
0
20
40
60
80
100
region (%)

B

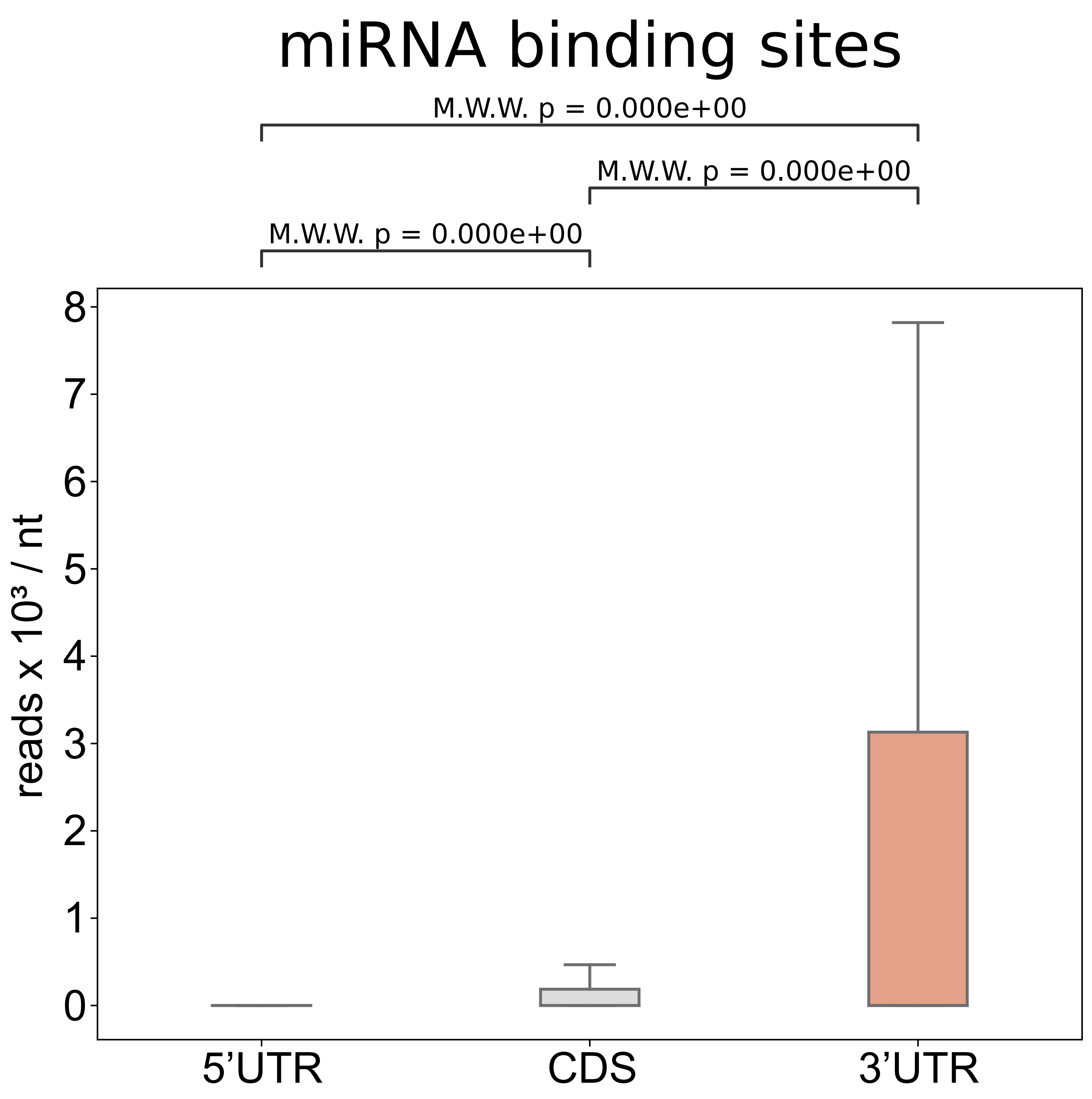
miRNA binding sites
M.W.W. p = 0.000e+00
M.W.W. p = 0.000e+00
M.W.W. p = 0.000e+00
reads x 10³ / nt
0
1
2
3
4
5
6
7
8
5'UTR
CDS
3'UTR

### Figure S2

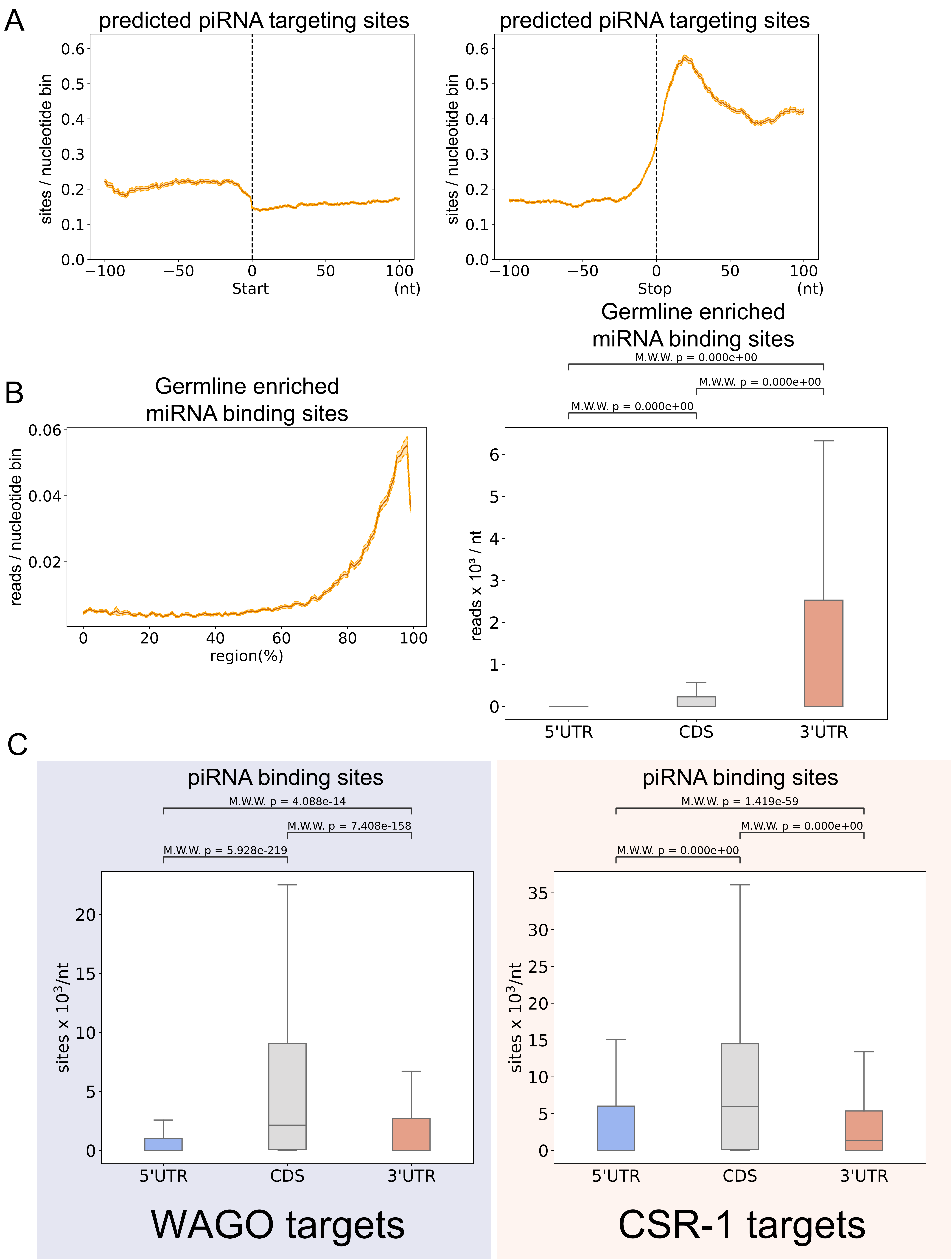

A
predicted piRNA targeting sites
sites / nucleotide bin
Start
(nt)
predicted piRNA targeting sites
sites / nucleotide bin
Stop
(nt)
B
Germline enriched miRNA binding sites
reads / nucleotide bin
region(%)
Germline enriched miRNA binding sites
M.W.W. p = 0.000e+00
M.W.W. p = 0.000e+00
M.W.W. p = 0.000e+00
reads x 10³ / nt
5'UTR
CDS
3'UTR
C
piRNA binding sites
M.W.W. p = 4.088e-14
M.W.W. p = 7.408e-158
M.W.W. p = 5.928e-219
sites x 10³/nt
5'UTR
CDS
3'UTR
WAGO targets
piRNA binding sites
M.W.W. p = 1.419e-59
M.W.W. p = 0.000e+00
M.W.W. p = 0.000e+00
sites x 10³/nt
5'UTR
CDS
3'UTR
CSR-1 targets

### Figure S3

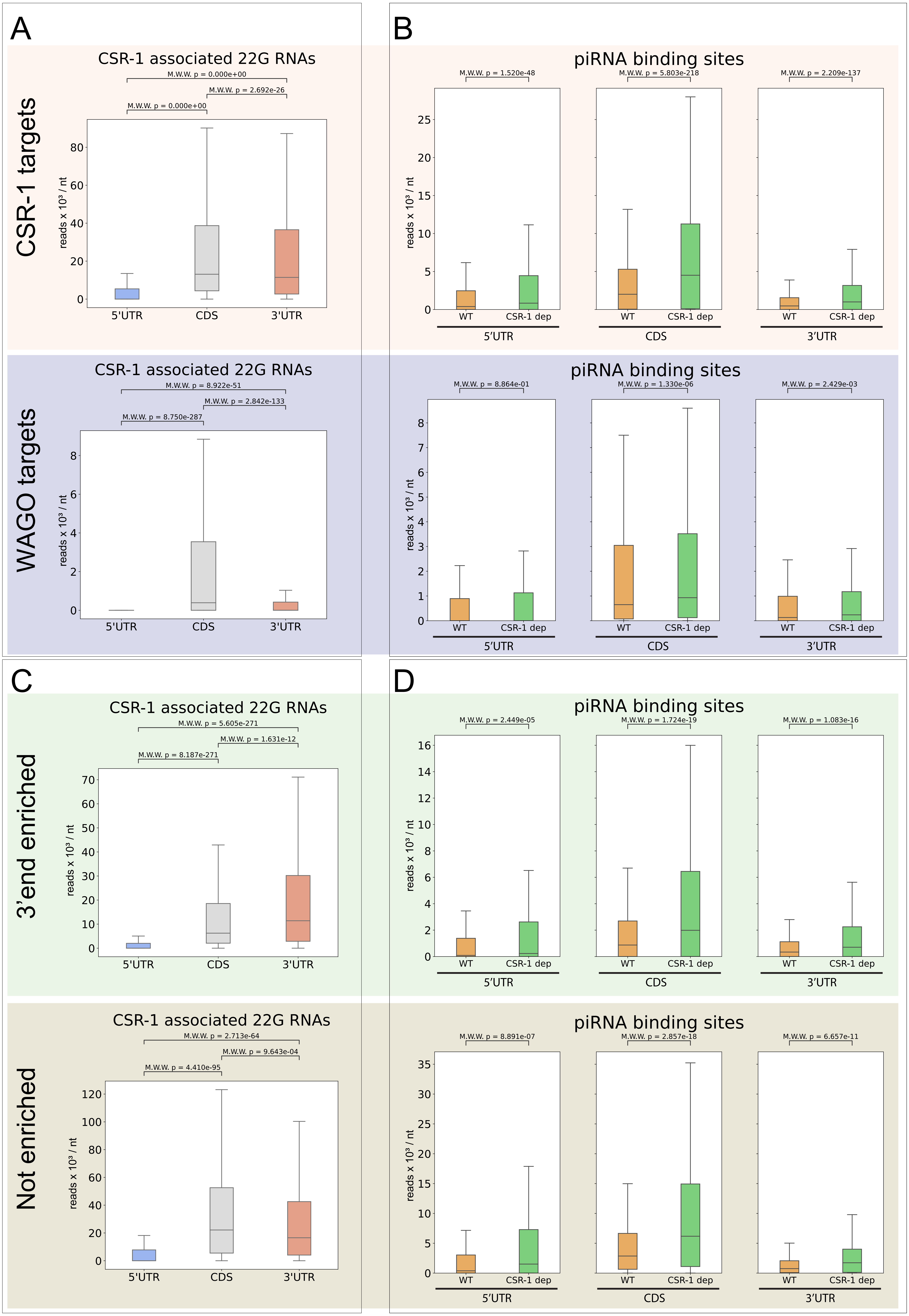

A
CSR-1 targets
CSR-1 associated 22G RNAs
M.W.W. p = 0.000e+00
M.W.W. p = 2.692e-26
M.W.W. p = 0.000e+00
reads x 10³ / nt
5'UTR
CDS
3'UTR
WAGO targets
CSR-1 associated 22G RNAs
M.W.W. p = 8.922e-51
M.W.W. p = 2.842e-133
M.W.W. p = 8.750e-287
B
piRNA binding sites
M.W.W. p = 1.520e-48
M.W.W. p = 5.803e-218
M.W.W. p = 2.209e-137
WT
CSR-1 dep
piRNA binding sites
M.W.W. p = 8.864e-01
M.W.W. p = 1.330e-06
M.W.W. p = 2.429e-03
C
3'end enriched
CSR-1 associated 22G RNAs
M.W.W. p = 5.605e-271
M.W.W. p = 1.631e-12
M.W.W. p = 8.187e-271
Not enriched
CSR-1 associated 22G RNAs
M.W.W. p = 2.713e-64
M.W.W. p = 9.643e-04
M.W.W. p = 4.410e-95
D
piRNA binding sites
M.W.W. p = 2.449e-05
M.W.W. p = 1.724e-19
M.W.W. p = 1.083e-16
piRNA binding sites
M.W.W. p = 8.891e-07
M.W.W. p = 2.857e-18
M.W.W. p = 6.657e-11

### Figure S3, continued

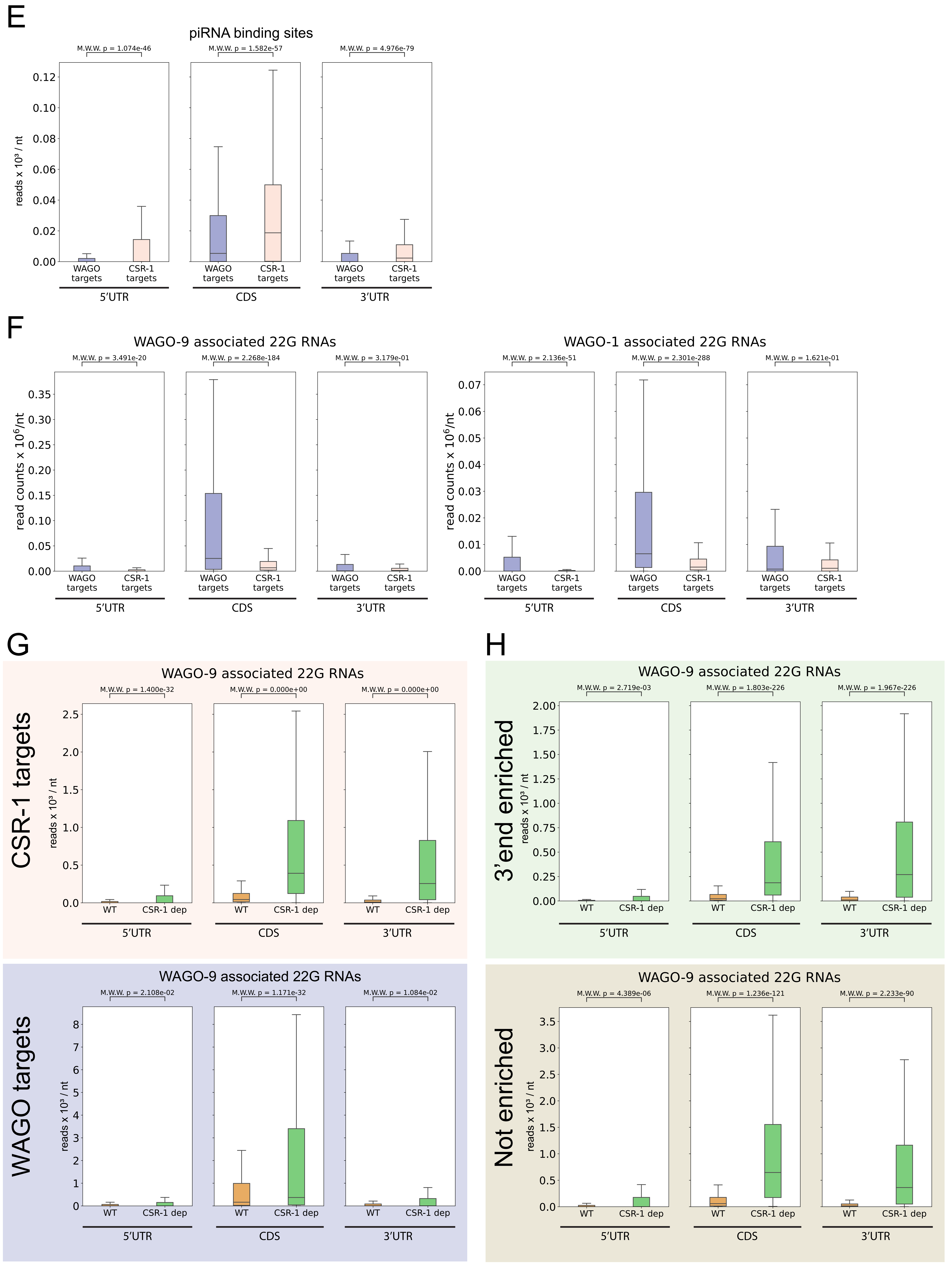

### Figure S4

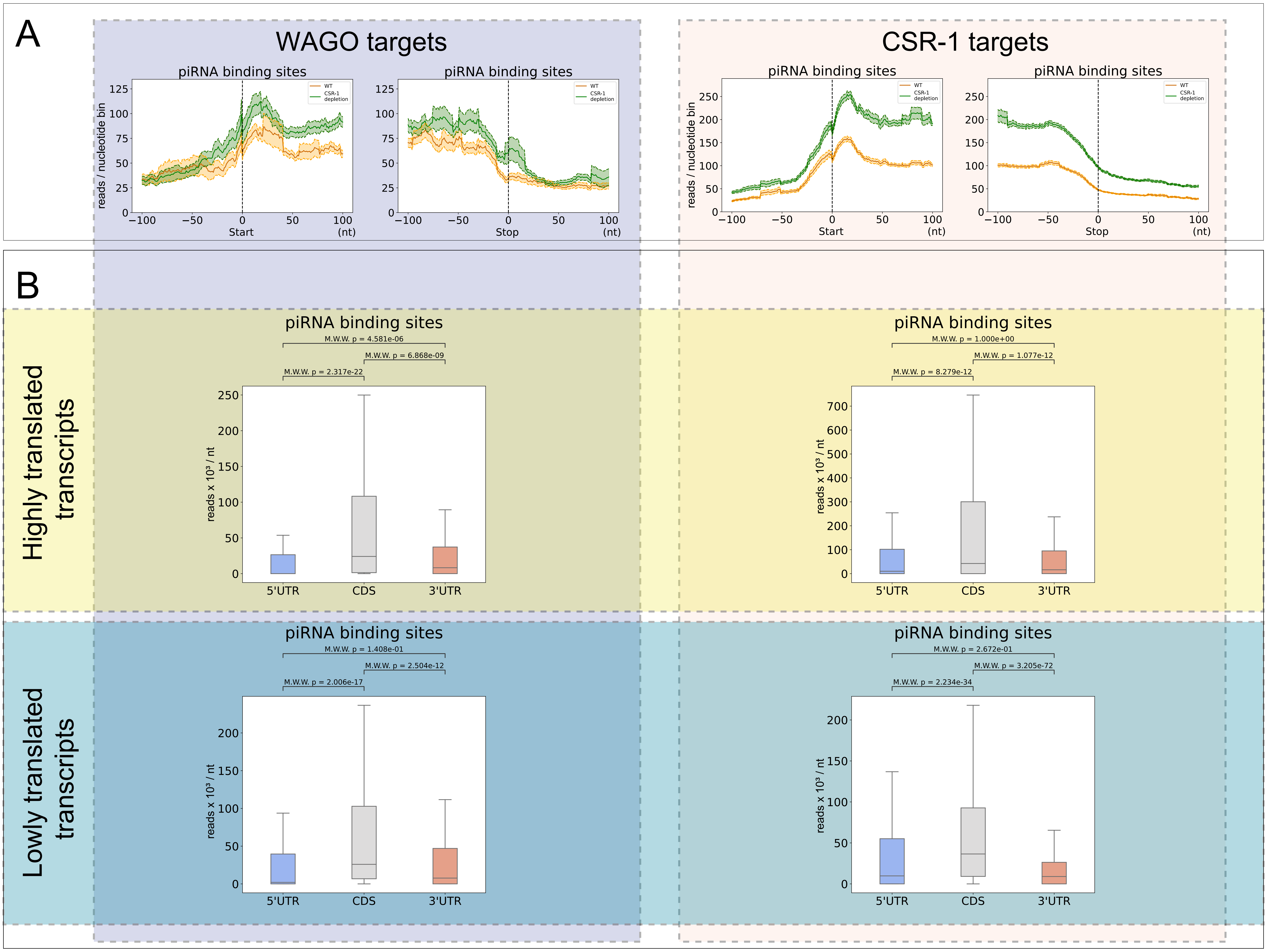

A
WAGO targets
CSR-1 targets
piRNA binding sites
reads / nucleotide bin
WT
CSR-1 depletion
Start
Stop
(nt)
B
Highly translated transcripts
Lowly translated transcripts
M.W.W. p = 4.581e-06
M.W.W. p = 6.868e-09
M.W.W. p = 2.317e-22
M.W.W. p = 1.000e+00
M.W.W. p = 1.077e-12
M.W.W. p = 8.279e-12
M.W.W. p = 1.408e-01
M.W.W. p = 2.504e-12
M.W.W. p = 2.006e-17
M.W.W. p = 2.672e-01
M.W.W. p = 3.205e-72
M.W.W. p = 2.234e-34
reads x 10³ / nt
5'UTR
CDS
3'UTR
